# Double-Strand Break Repair Pathways Differentially Affect Processing and Transduction by Dual AAV Vectors

**DOI:** 10.1101/2023.09.19.558438

**Authors:** Anna C. Maurer, Brian Benyamini, Vinson B. Fan, Oscar N. Whitney, Gina M. Dailey, Xavier Darzacq, Matthew D. Weitzman, Robert Tjian

**Affiliations:** Department of Molecular and Cell Biology, University of California, Berkeley, CA, USA; CIRM Center of Excellence, University of California, Berkeley, CA; Li Ka Shing Center for Biomedical & Health Sciences, University of California, Berkeley, CA, USA; University of Pennsylvania Perelman School of Medicine and the Children’s Hospital of Philadelphia, Philadelphia, PA, USA; Howard Hughes Medical Institute, University of California, Berkeley, CA, USA

## Abstract

Recombinant adeno-associated viral vectors (rAAV) are a powerful tool for gene delivery but have a limited DNA carrying capacity. Efforts to expand this genetic payload have focused on engineering the vector components, such as dual trans-splicing vectors which double the delivery size by exploiting the natural concatenation of rAAV genomes in host nuclei. We hypothesized that inefficient dual vector transduction could be improved by modulating host factors which affect concatenation. Since factors mediating concatenation are not well defined, we performed a genome-wide screen to identify host cell regulators. We discovered that Homologous Recombination (HR) is inhibitory to dual vector transduction. We demonstrate that depletion or inhibition of HR factors BRCA1 and Rad51 significantly increase reconstitution of a large split transgene by increasing both concatenation and expression from rAAVs. Our results define new roles for DNA damage repair in rAAV transduction and highlight the potential for pharmacological intervention to increase genetic payload of rAAV vectors.

## INTRODUCTION

Recombinant adeno-associated viral vectors (rAAV) are a leading platform for clinical DNA delivery. This vector system is derived from a human parvovirus and has many inherent features that make it attractive for human gene therapy applications^1^. It also has some inherent limitations, the major one of which is the small genetic payload. The unenveloped 25 nm capsid can package no more than 5 kb of single-stranded DNA (ssDNA) into its small lumen^2,3^, imposing a size limit to the coding and regulatory elements that can be included in the transgene cassette. For gene replacement therapies, current strategies employ cDNA versions of the gene of interest, with expression driven by a short, strong promoter such as CMV^4^. While this accommodates many therapeutically relevant gene sizes, it excludes other larger genes, such as dystrophin or the most accurate Cas9 versions and other base editors. Moreover, CMV and similar promoters are ubiquitously expressed at very high levels, leading to off-target expression in non-therapeutically relevant tissues, and potentially toxic levels of transgene expression. This can lead to immune responses and other failures of the therapy. A larger payload size is therefore highly desirable for expressing large proteins and moreover for enabling fine-tuned spatiotemporal control over expression and levels by inclusion of more diverse, efficient regulatory sequences.

The wild-type (wt) AAV virus and the recombinant AAV vector have the same overall structure, consisting only of the 60mer icosahedral capsid with one packaged ssDNA vector genome (VG). Viral proteins are required during production for replication and packaging, but no viral enzymes are packaged within the virion^5^. VG architecture requires flanking the desired delivery DNA with the viral inverted terminal repeat (ITR) sequences at both ends^6^. The ITR contains sequence elements that are required during viral replication and also direct packaging into the preformed capsids for vector production^7^. In the transduction process (Figure 1A), the capsid traffics the VG to the host/target cell nucleus, where the capsid is shed (termed uncoating) ^8,9^. The ssDNA VG is converted to dsDNA through unclear mechanisms which enables transgene expression. Individual VGs can also circularize through intramolecular ITR-ITR joining and concatenate through intermolecular ITR-ITR joining^10^. In its recombinant form, where a capsid of choice is packaged with an ITR-flanked transgene, rAAV VGs cannot replicate; but instead multiple VGs enter the nucleus and can concatenate into large episomal species^11^. The absence of delivered enzymes implies that the post-uncoating nuclear steps of rAAV transduction are orchestrated by host cellular factors. Decades of observation have articulated the nuclear steps of transduction – uncoating, conversion to dsDNA, concatenation, expression – yet they remain observations, with little mechanistic detail and knowledge of facilitating host factors^12,13^

**Figure 1.**
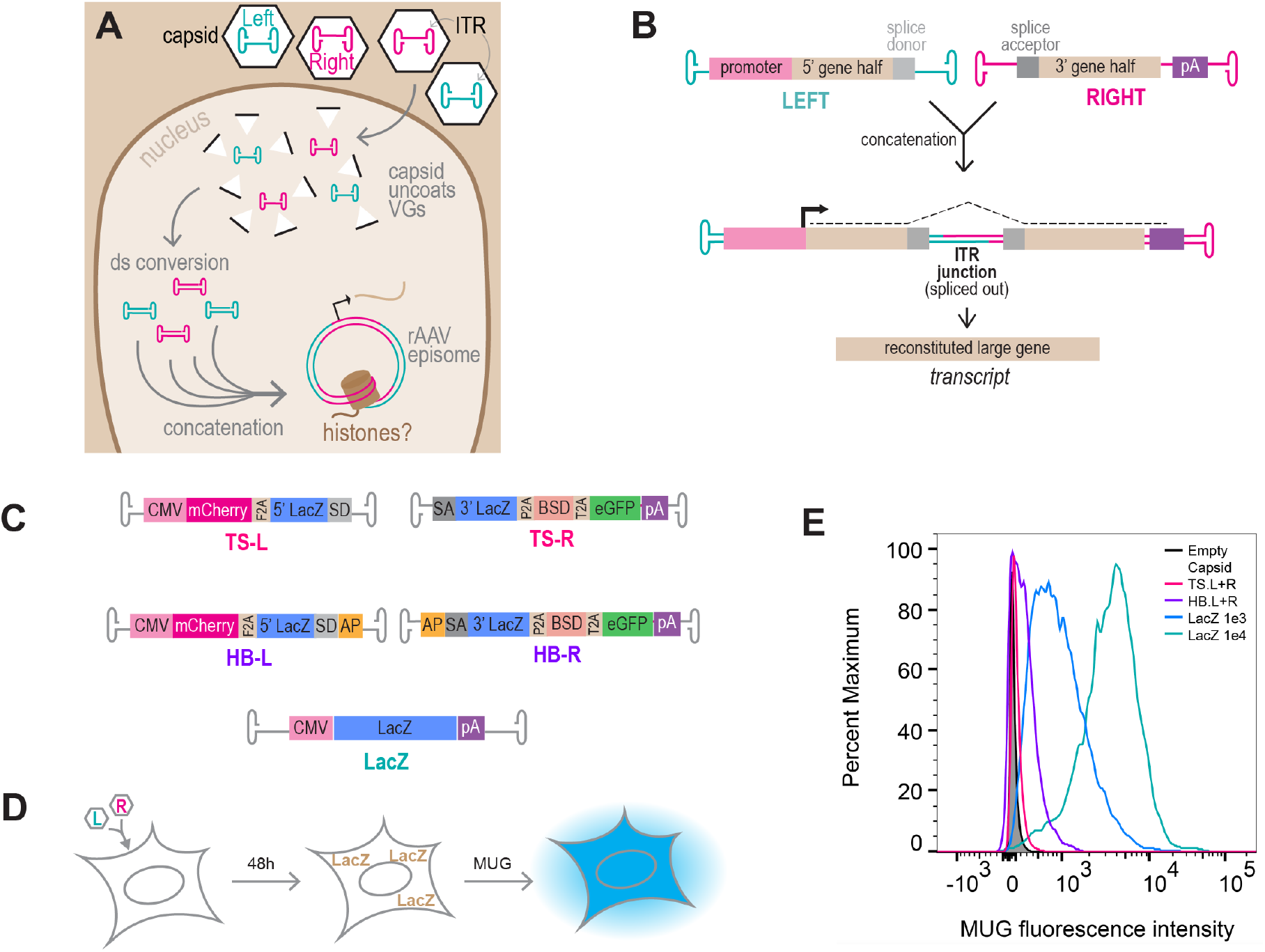
Dual vectors are less efficient than an unsplit transgene. **(A)** Schematic of rAAV vector genome (VG) processing. ssDNA VGs are converted to dsDNA and VGs circularize and concatenate by ITR-ITR joining, through unclear mechanisms. VGs may associate with histones, but at what point during these processes is unknown. **(B)** The dual vector approach can double payload size by exploiting the natural concatenation process. A large gene is split into separate Left (L) and Right (R) VGs; inclusion of a splice donor/acceptor excludes the ITR junction from the final transcript. **(C)** Reporter constructs to assay split transgene reconstitution. LacZ is split between L and R VGs in a trans-splicing (TS) pair that requires ITR-ITR concatenation for expression, or a Hybrid (HB) trans-splicing which additionally includes the Alkaline Phosphatase (AP) partial sequence previously demonstrated to increase split transgene reconstitution efficiency. An unsplit LacZ VG is used as a control. **(D)** LacZ Activated Fluorescence Assay (LAFA) schematic: cultured cells are transduced with equimolar doses of L and R vectors. After 48h, split transgene reconstitution is assayed by Beta-galactosidase (LacZ) activity cleaving 4MU beta-D-galactopyranoside (MUG), a fluorogenic beta galactose analog. **(E)** U2-OS cells were treated with 1e4 VG/cell of each L and R vectors (shown in red for TS and purple for HB), unsplit vector (shown in blue and green for two different doses), or empty capsid (shown in black) negative control. LAFA was performed at 48 hours post transduction and read by flow cytometry. All capsids and ITRs are based on AAV2.

The absence of viral enzymes in the rAAV particle implies that the processes of VG circularization and concatenation are orchestrated by host cellular factors. ITR-ITR joining is analogous to host DNA repair mechanisms, and dsDNA VGs could conceivably resemble a chromosomal break. The host cell has many ways of repairing DNA damage, including two main pathways to repair double-strand breaks (DSB): the high-fidelity but restricted activity pathway called Homologous Recombination (HR) or Homology Directed Repair (HDR), and the constitutively active but error-prone pathway of Non-Homologous End Joining (NHEJ)^14^. The MRE11/RAD50/NBS1 (MRN) complex, which recognizes free DNA ends to facilitate repair upstream of HDR versus NHEJ pathway choice, recognizes ITRs but inhibits concatenation^15,16^. The NHEJ pathway has been suggested to play a role in concatenation since depletion or inhibition of the NHEJ factor DNA-PKcs reduces VG concatenation^17–21^. However, the effect of HDR pathway on concatenation has not been directly examined.

The propensity of VGs to concatenate can be exploited to increase the final genetic payload size^22^. In this “dual vector” approach (Figure 1A-B), a large transgene cassette is split into two halves, packaged into separate capsids, and co-delivered to target cells. Inclusion of a splice donor in the “Left” half VG (L) and a splice acceptor in the “Right” half VG (R) allows the ITR junction to be spliced out during RNA processing of transcription, and the large gene is subsequently reconstituted in the transcript. This technique was first developed in the early 2000s by multiple groups independently^23–25^. To be effective it requires correct, directional, and stoichiometric concatenation, followed by correct expression. By nature of the sequence, any ITR can in theory join any ITR, leading to many permutations that do not all lead to correct split transgene reconstitution. The approach also currently relies on very high vector doses. To increase the chances of L and R genomes joining in the correct orientation, a short fragment of the Alkaline Phosphatase (AP) gene serendipitously found to be “highly recombinogenic” in the dual vector context^26^ can be added between the splice signal and the ITR^27^. It is presumed that the AP fragment promotes annealing of + and – sense VGs which then become a reconstituted large gene through HR. However, HR has never been directly tested as the mechanism of concatenation. Indeed, AP inclusion significantly increases efficiency and is now commonly used in dual vectors, but like general mechanisms of concatenation, little is known about the mechanisms underpinning the AP phenomenon.

Here, we aimed to improve dual vector transduction by an alternative approach that focuses on the host cell. We first identify cellular pathways inhibitory to split transgene reconstitution by a genome-wide screen. We provide the first evidence for early chromatinization of VGs, and epigenetic recognition as a double-strand break. Surprisingly, blocking HR increases both concatenation of and expression from rAAV VGs, which dramatically increases dual vector transduction efficiencies. Our findings suggest that HR is the “first responder” to the DSB-flagged VGs, but that NHEJ is a more efficient mechanism for VG concatenation. While the majority of efforts to improve rAAV gene therapies have focused on modifying the vector and dose, our findings emphasize the importance of understanding host responses in the target cell. Our results highlight the potential for pharmacological approaches to enhance efficiency and increase deliverable transgene size in rAAV gene therapies.

## RESULTS

### A genome-wide screen to identify cellular inhibitors of split transgene reconstitution

To identify cellular factors whose modulation could increase dual vector transduction, we established a platform amenable to pooled genome-wide screens by building upon dual vector systems previously engineered by others in which vectors enable trans-splicing (TS) or contain the additional AP segments in Hybrid (HB) vectors ^23,25,24,28,29,27^ (Figure 1C). Correct directional concatenation and successful expression will yield a multi-cistronic cassette containing several reporter genes. The LacZ gene is split to span the ITR junction between Left (L) and right (R) vectors and is therefore the most stringent reporter of concatenation and subsequent expression. After transducing cultured cells, incubation with the fluorogenic B-galactose analog 4-methylumbelliferyl-[beta]-D-galactopyranoside (MUG) provides an enzymatic readout of successful split transgene reconstitution. This LacZ activated fluorescence assay (LAFA) can be performed on cell lysates in microwell plates or in flow cytometry based approaches and enables FACS isolation of transduced cells (Figure 1D). Flow cytometry of treated cells (Figure 1E) confirmed that at 1e4 total vg/cell (5e3 vg/cell of L vector plus 5e3 vg/cell of R vector) for both the TS (magenta) and HB (purple) dual vectors transduced at efficiencies two orders of magnitude lower than cells receiving the same dose of an unsplit LacZ vector (green). The inefficiency of dual vector transduction is further demonstrated by comparing 1e4 total VG/cell dual vectors (purple) to 1e3 VG/cell unsplit LacZ (blue); a 10-fold lower dose of unsplit transgene achieves 10-fold higher transduction than dual vectors.

To identify opportunities for host factor modulation that could lead to increases in split transgene reconstitution, we reasoned that (1) most pharmacological intervention presents inhibitors rather than activators, and (2) as a virally derived entity, it is likely that rAAV concatenation and expression are targets of negative regulation as part of an innate defense. We therefore designed a genome-wide knockout (KO) screen to identify repressors of dual vector transduction. Lentiviral Brunello libraries carrying Puromycin resistance cassette, Cas9, and an average of four 4 guides per gene were used to generate genome-wide KO libraries of U2-OS cells. After Puro selection, the cellular KO libraries were co-transduced with 5e3 vg/cell of each of the HB-L and HB-R vectors. Cells were collected at 48 hours post-transduction (hpt), stained with MUG, and LacZ expressing cells isolated by FACS. Genetic perturbations in these cells were identified by NGS, and statistical analysis on guide representation in this pool was performed to generate a list of candidate inhibitors of dual vector transduction (Figure 2A) Pathway analysis of the top 2% of significantly enriched genes in the dual vector transduced population revealed overrepresentation in DNA synthesis and repair, particularly double-strand break (DSB) repair. Of the two major DSB repair pathways, Homology Directed Repair (HDR) and Non-Homologous End Joining (NHEJ), only HDR was overrepresented (Figure 2B). Since the NHEJ pathway has been suggested to regulate concatenation in a positive way, we hypothesized that the HDR DSB repair pathway could function as a possible negative regulator of VG concatenation. This was addressed in our subsequent hit validation.

**Figure 2.**
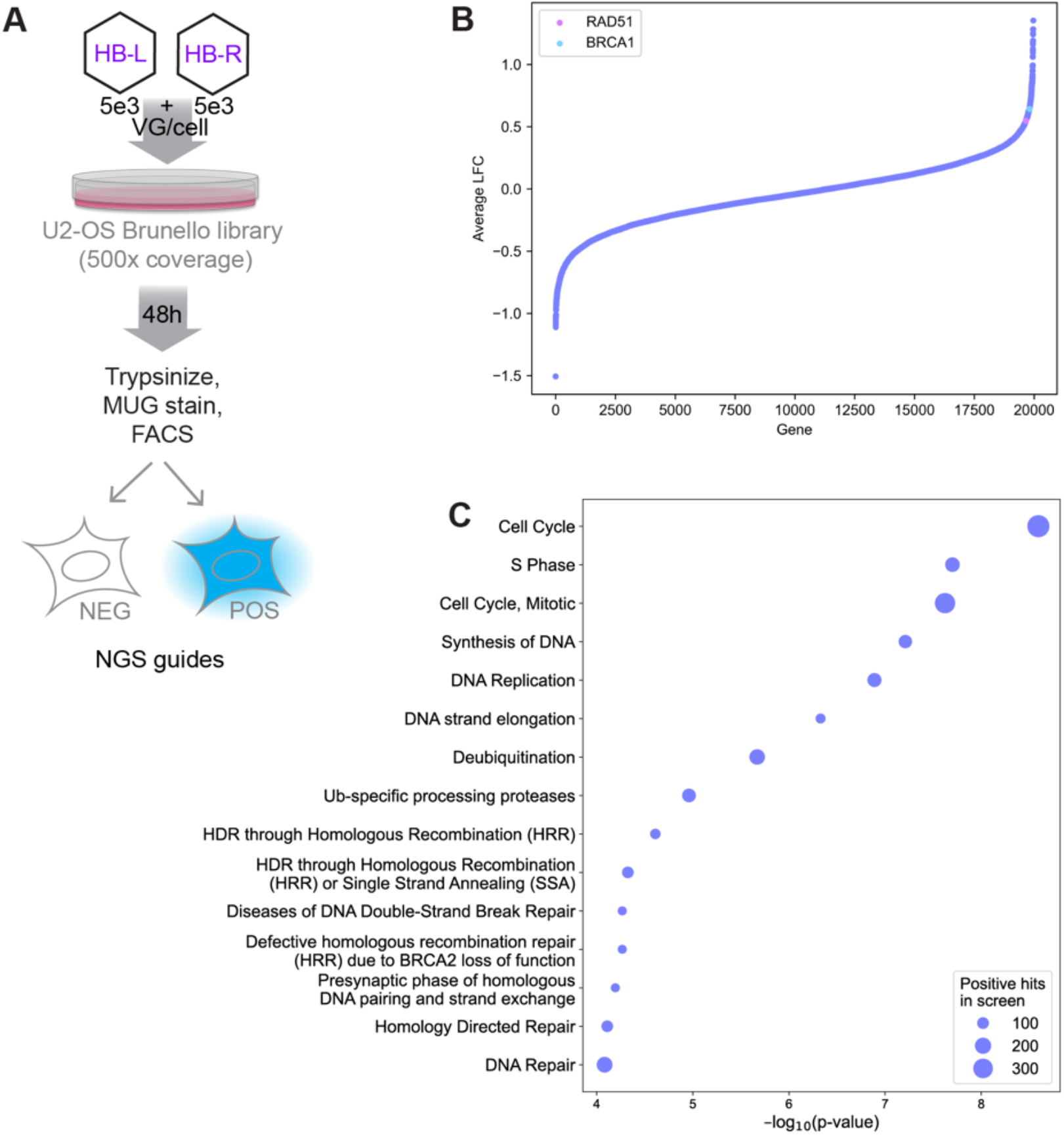
A genome-wide screen to identify cellular inhibitors of split transgene reconstitution. (A) Screen schematic: a Brunello KO Library of U2-OS cells (500x coverage) was treated with 5e3 vg/cell of the HB dual vectors. Cells were collected at 48 hpt, stained with MUG, and sorted by FACS. (B) Average fold-change of guide abundance (positive vs negative population) sorted and plotted as a waterfall plot. Rad51 and BRCA1 are highlighted. (C) Reactome pathway analysis results on the top 5% of genes in the positive population (candidate negative regulators of split transgene reconstitution), sorted by significance. Bubble size indicates the number of genes in each Reactome category.

### Vector genomes are epigenetically marked as double-strand breaks

Since chromosomal DSBs are marked by phosphorylation of the histone variant H2AX (gH2AX) which coordinates recruitment of repair factors, we examined whether VGs are subject to this epigenetic mark. Previous studies have observed an increase in nuclear gH2AX when cells are infected with wt AAV^30–32^, but this epigenetic mark has not yet been examined on AAV viral or vector genomes themselves. To detect individual VGs, we adapted a previously established approach^15,33–36^ in which the rAAV genome is visually detected through recognition of *lacO* binding sites by a fluorescent LacI fusion protein. We generated U2-OS cells stably expressing the LacI-mNeonGreen fusion protein, and transduced with rAAV vectors containing VGs consisting of 64 LacO repeats. After uncoating and conversion to a dsDNA species, the VG recruits many fluorophores and can be visualized in live or fixed cells (Figure 3A). We observed VG foci distributed throughout the nucleus, fusing with other foci, and increasing in size and intensity over time, potentially reflecting concatenation events (Movie 1). We first asked whether VG foci are marked as DSBs by immunostaining for gH2AX over a 4-48 h time course. We observed colocalization as early as 4 hpt (Figure 3B). To date, this is the earliest observed histone association with rAAV-delivered DNA and suggests that VGs are chromatinized and flagged as DSBs quickly after uncoating.

**Figure 3.**
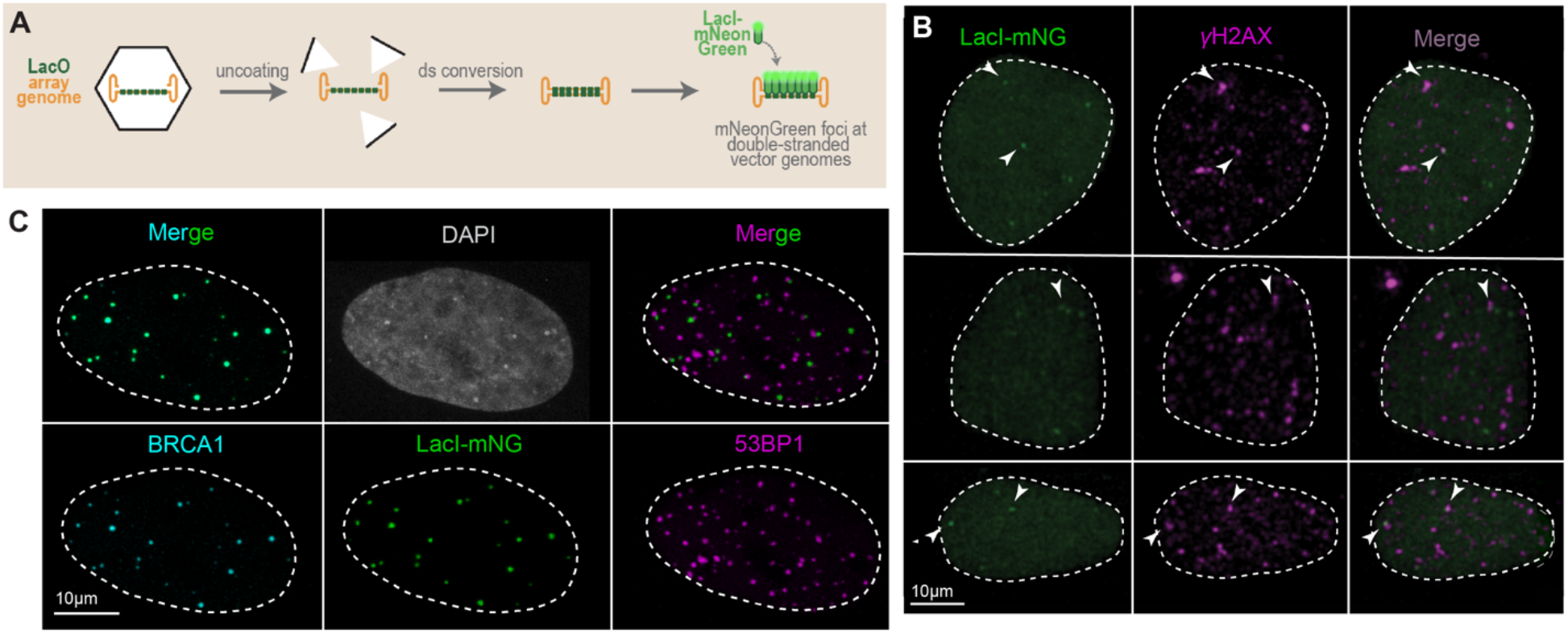
rAAV vector genomes (VG) are recognized as double strand breaks (DSB). **(A)** VG visualization schematic. Capsids are packaged with array VGs comprised of 64 lacO repeats. After uncoating and becoming double-stranded (dsDNA) in LacI-mNeonGreen expressing cells (U2-OS^LacI-mNeonGreen^), VGs can be visualized as green foci. U2-OS^LacI-mNeonGreen^ cells were treated with 1e5 VG/cell of AAV2/2.LacO.64, fixed at **(B)** 4 hpt and immunostained for yH2AX, or **(C)** 48 hpt and immunostained for BRCA1 and 53BP1.

We next asked whether VGs recruit HDR and NHEJ factors by immunostaining. We used antibodies to BRCA1 as a representative HR factor which was enriched in our screen, and antibodies to 53BP1 as a representative NHEJ factor. Colocalization of both repair factors were observed with VG foci, however we frequently observed complete overlap of VG with BRCA1 foci, whereas 53BP1 foci were less frequently overlapping VG foci (Figure 3). These anti-correlated staining patterns are in line with our opposing screen results for NHEJ and HDR, and the known competition between these DNA repair pathways on DSBs in the cellular genome^14^.

### HDR deficiency increases concatenation and expression from rAAV vectors

Successful trans-splicing dual vector transduction requires two major steps: concatenation of the L and R co-infected VGs followed, by successful expression of the split cassette. To examine whether our candidate factors affect concatenation or expression, we added an mScarlet expression cassette downstream of the lacO array (lacO.64.CMV.mScarlet; Figure 4A). This enables simultaneous and quantitative visual readouts of both concatenation (mNeonGreen foci) and expression (mScarlet intensity) which are not interdependent, such as with dual vector transduction and LacZ reconstitution. To test effects of BRCA1 loss on concatenation and expression, we introduced BRCA1 or scrambled guides by transfection into U2-OS^LacI-mNG,Cas9^ cells, waited 72h for editing, and then transduced with rAAV2/2.lacO.64.CMV.mScarlet vectors. BRCA1 levels, VG foci and mScarlet expression were quantified in thousands of cells by high-content imaging at 48 hpt. We observed significant increases in the average number of VG foci per cell (Figure 4B) and the percentage of cells with any VG foci in cells receiving BRCA1 guides compared to Scramble (Figure 4C). We next quantified expression by mScarlet intensity in the high-content images. Expression of the mScarlet transgene is 10% higher in cells receiving BRCA1 guides, a small but statistically significant increase (Figure 4D). Together these observations suggest that BRCA1 inhibits concatenation and *de novo* episome formation, which impacts gene expression.

**Figure 4.**
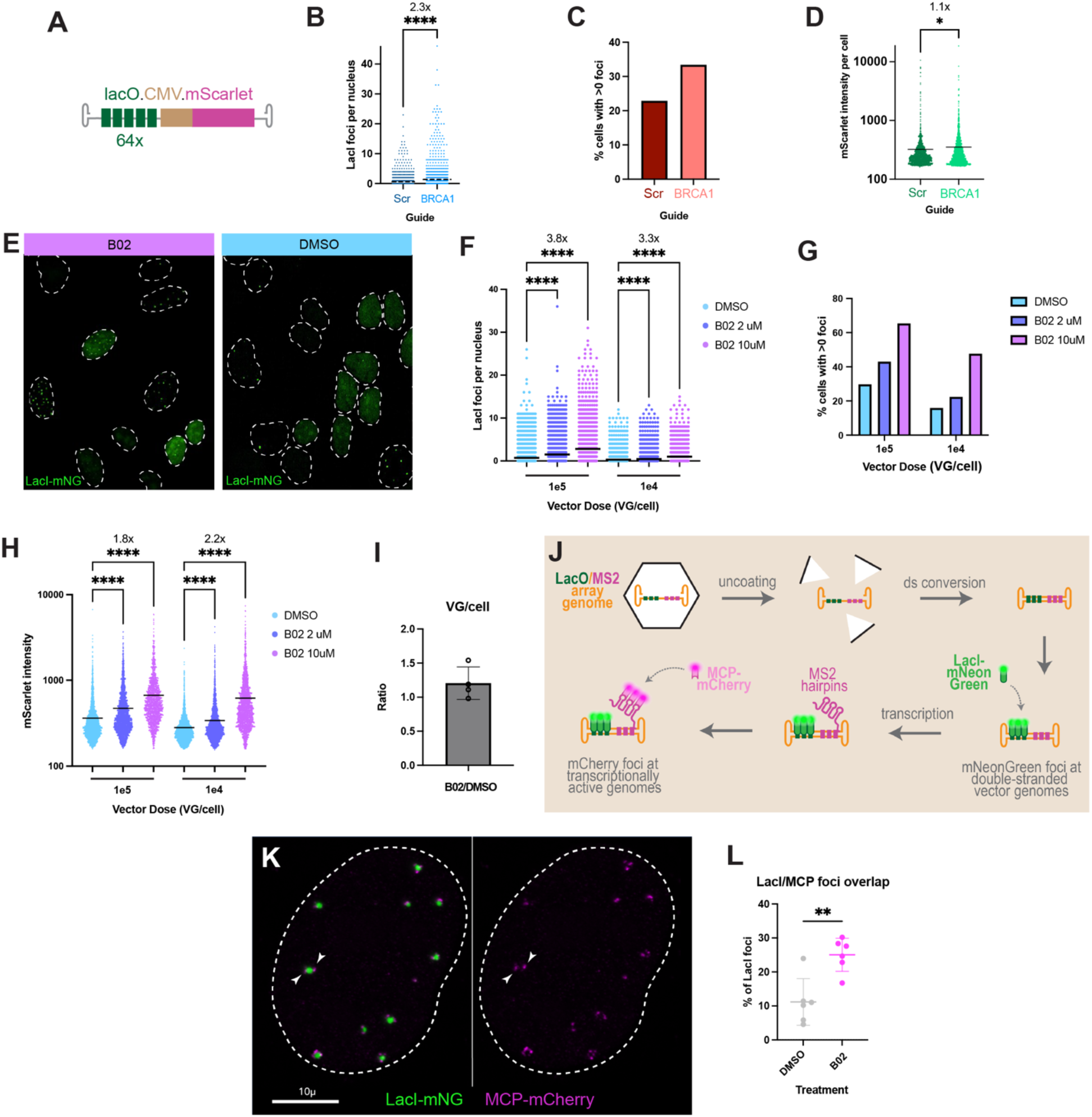
HDR deficiency increases concatenation and expression from rAAVs. U2-OS^LacI-mNeonGreen^ cells stably expressing Cas9 were transfected with BRCA1 or Scramble guide expressing plasmids, placed under selection for 72h, then transduced with 1e5 vg/cell of **(A)** AAV2/2.lacO.CMV.mScarlet and high-content imaged 48 hpt **(B-D**). Individual cells are plotted as points according to y axis parameter, or as a bar graph for percentage parameters. Black lines = mean value; fold increase indicated above graph. **(E-H)** U2-OS^LacI-mNeonGreen^ cells were pretreated with drug or DMSO (legend), transduced with AAV2/2.lacO.CMV.mScarlet at dose on x-axis, then high-content imaged at 48 hpt. **(E)** Primary image example. **(F)** Individual cells are plotted as points by number of foci per nucleus. Fold increase indicated above graph; black line = mean. **(G)** Percentage of all cells in (F) with any foci (>0). **(H)** Images were segmented by cell and transgene expression levels assessed by average mScarlet pixel intensity, then plotted as individual points. Fold increase indicated above graph; black line = mean. **(I)** Vector copy number per cell determined by qPCR 48 hpt with 1e5 vg/cell, in DMSO vs B02 treated cells. n=4 biological replicate wells are plotted as individual points as the ratio of B02:DMSO copy numbers; error bar = SD. Statistical significance was determined by a paired, two-tailed T test (P=0.1599) **(J)** Schematic of lacO.MS2 system to quantify transcriptionally active VGs. **(K)** U2-OS^LacI-mNeonGreen, MCP-mCherry^ cells were treated with 1e5 VG/cell AAV2/2.lacO.CMV.MS2 and imaged 48hpi on an LSM900 Airyscan2. Arrowheads indicate multiple actively transcribing VGs in a single episome. **(L)** U2-OS^LacI-mNeonGreen, MCP-mCherry^ cells were treated with 1e5 VG/cell AAV2/2.lacO.CMV.MS2 and high-content imaged at 48 hpt. The percentage of LacI foci with at least one overlapping MCP focus in triplicate wells and in biological duplicate are plotted as individual points. Statistical significance was determined by unpaired, two-tailed T tests in (A), (D), and (L), and by Kruskal-Wallis Test with Dunn’s multiple comparisons test in (F) and (H), with *p≤0.05; **p≤0.01; ***p≤0.001; ****p≤0.0001.

To interrogate whether the increased foci formation is specific to BRCA1 loss or is an effect of non-functional HDR, we tested the effects of Rad51 inhibition as another HDR factor enriched in our screen. Aiming for a more complete penetrance and an approach that is potentially translatable to clinical use, we inhibited Rad51 with the drug B02 by pretreating U2-OS^LacI-mNG^ cells with multiple doses of drug, then transduced with multiple doses of rAAV2/2.lacO.64.CMV.mScarlet. We observed that B02 increased the number of VG foci per cell (Figure 4E, F) and the percentage of cells forming any VG foci (Figure 4G), in a dose-dependent fashion and exceeded the effects of BRCA1 loss. Additionally, high-dose Rad51 inhibitor resulted in approximately doubled mScarlet expression, whether vector was at moderate (1e4 VG/cell) or high doses (1e5 VG/cell) (Figure 4H). These results suggest that, similar to BRCA1, Rad51 inhibits concatenation, *de novo* episome formation, and transgene expression. Together these results implicate repressive roles for HDR in rAAV transduction.

To examine whether the observed increases in VG foci and transgene expression were due to increased total number of VGs present in cells, for example due to increased nuclear import or increased dsDNA conversion leading to increased stability/decreased degradation in HDR deficient cells rather than increased concatenation or expression activity, we compared VG copy number in DMSO and B02 treated cells 48 hpt. Cells treated with B02 have no significant difference in VG copy number compared to cells treated with DMSO and receiving the same vector dose (Figure 4I). Taken together, these observations confirm that HDR inhibits both concatenation and expression of rAAV VGs.

### Rad51 inhibition increases the proportion of transcriptionally active vector genomes

We next asked whether the increase in expression we observe under Rad51 inhibition is a result of increased transcriptional activity from each active VG or an increase in the proportion of VGs that are transcriptionally active. To observe transcriptional activity *in situ* with each VG, we replaced the mScarlet gene with an MS2 array which forms stem-loops when transcribed. In MCP-mCherry expressing cells the nascent transcript can be visualized as an mCherry focus overlapping with an mNeonGreen (VG) focus (Figure 4J). Transducing U2-OS^LacI-mNG, MCP-mCherry^ cells with rAAV2/2.lacO.64.CMV.MS2 array vectors, we observed large VG foci often with multiple actively transcribing units within the concatemer (Figure 4K). After pretreatment with B02 or DMSO, high-content imaging at 48 hpt revealed a significant increase in the percentage of VG foci with at least one overlapping MS2 focus in B02 treated cells (Figure 4L). This suggests that Rad51 inhibition increases expression by increasing the proportion of transcriptionally active genomes.

Given the inhibitory role we observed for HDR in concatenation and expression independently, we next tested the effects of B02 on dual vector transduction, which requires both processes. We pretreated cells with multiple doses of B02, transduced cells with multiple doses of the Trans-Splicing (TS) L+R split LacZ vector (Figure 1C), and used LAFA (Figure 1D) in a 96-well format to readout split transgene reconstitution at 48 hpt (Figure 5A). At all dose combinations, B02 significantly increased dual vector expression. The most dramatic effect was seen at the lowest vector doses, at which 10 mM B02 increased split LacZ expression 27-fold over DMSO treated cells receiving the same vector dose. This B02-induced increase in expression at low vector doses is comparable to expression levels achieved by 100-fold higher vector doses not treated with B02 (Figure 5A, grey dashed line). This suggests that chemically-induced improvements in transduction will enable significantly lower doses of vector to be used for gene replacement approaches.

**Figure 5.**
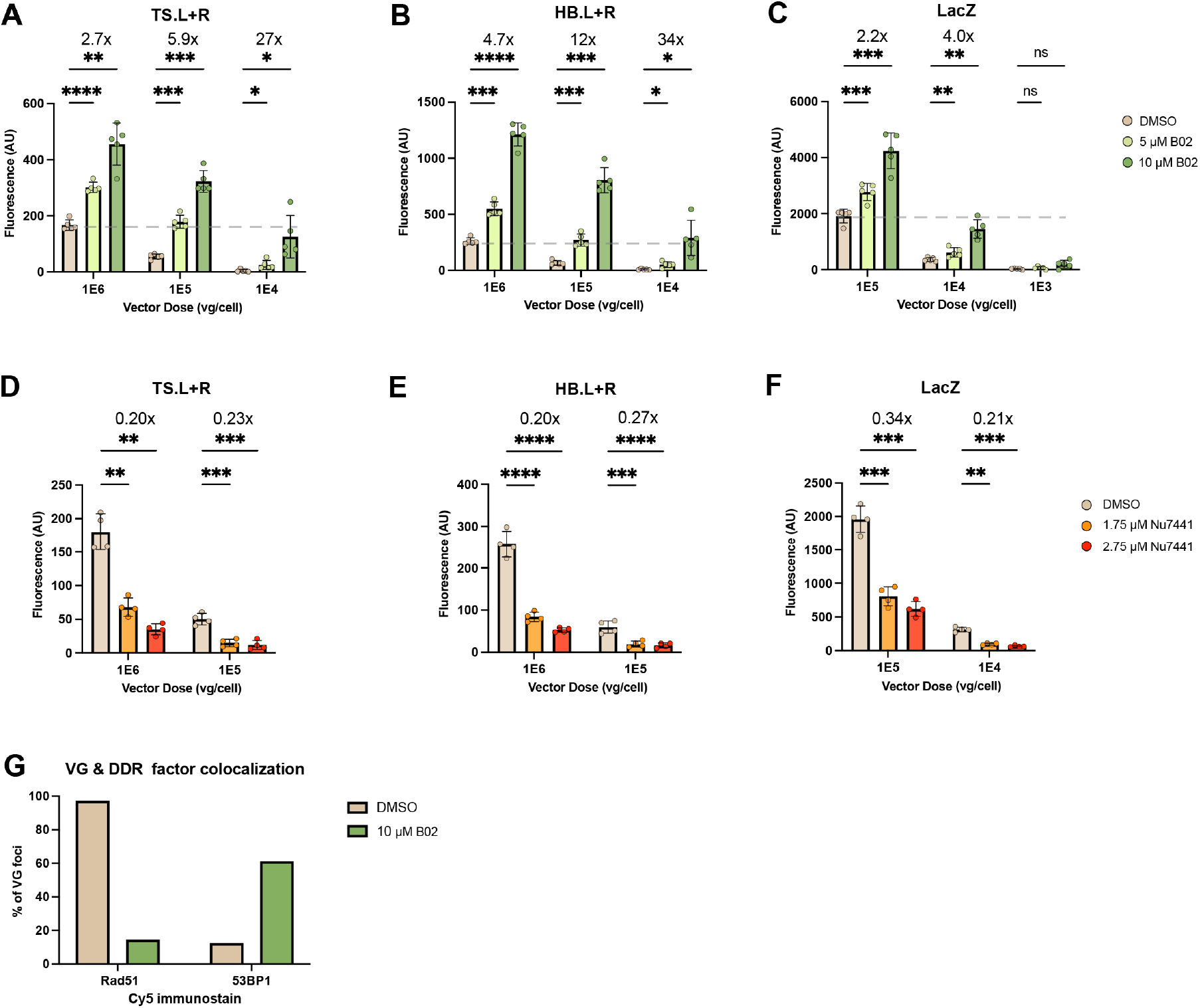
Rad51 inhibition increases dual vector transduction. The effects of **(A-C)** Rad51 or **(D-F)** DNA-PKcs inhibition on split transgene reconstitution were tested in U2-OS cells by LAFA 48 hours post-transduction by the dual (or single) vector indicated above each plot. Bars represent the mean of 5 **(A-C)** or 4 **(D-F)** biological replicates; individual values are plotted as dots, bars represent SD. A 2-way ANOVA was used, with *p≤0.05; **p≤0.01; ***p≤0.001; ****p≤0.0001. **(G)** U2-OSlacI-mNeonGreen cells were pretreated with DMSO or B02, transduced with 1e5 VG/cell AAV2/2.lacO.64, and immunostained for Rad51 (HDR) and 53BP1 (NHEJ) using a Cy5 secondary antibody at 48 hpt. VG and Cy5 foci overlap was quantified by high content imaging.

### The AP fragment increases concatenation by an HR-independent pathway

While the TS dual vectors serve to model the natural concatenation process and best represent our high-content imaging findings, our original screen was performed with the more efficient hybrid (HB) dual vectors (Figure 1C), for reasons of experimental practicality. Originally developed by Ghosh and colleagues^27^, this pair contains a “highly recombinogenic” overlapping sequence fragment from the alkaline phosphatase (AP) gene that bridges L and R vectors, providing an additional mode of reconstitution beyond the ITR-ITR concatenation in the traditional TS dual vectors (Figure 1B & C). The mechanism through which AP increases efficiency of dual vector transduction is not well understood; there are many mentions in the literature that the AP overlapping sequence drives split transgene reconstitution through homologous recombination, however HR or HDR have not yet been directly examined in this context. Surprisingly, B02 had even more potent effects increasing HB dual vector transduction than TS (Figure 5B), suggesting that the classical HR pathway is not the main mechanism through which hybrid dual vectors reconstitute a split transgene. Rad51 inhibition also increased unsplit LacZ vector transduction (Figure 5C), but to a much lower degree than dual vectors; at 1e4 VG/cell, unsplit LacZ expression is increased 4 fold by high dose B02, whereas L+R vectors at the same dose are increased up to 34 fold wit B02 (Figure 5A-C). This further corroborates our findings that HDR loss increases expression from VGs, and suggests that the concomitant increase in concatenation synergistically contributes to a dramatic increase in dual vector transduction.

The observation that HDR inhibits VG concatenation is counterintuitive, since concatenation could be considered as analogous to repairing a chromosomal DSB. HDR is a less active repair mechanism than its error-prone counterpart Non-Homologous End Joining (NHEJ), which coordinates DSB repair through factors distinct from HDR including 53BP1 and the DNA-dependent protein kinase catalytic subunit (DNA-PKcs). Previous studies recovered fewer circularized VGs from transduced muscle in DNA-PKcs deficient mice than in controls^18,37^, suggesting an important role for NHEJ in VG concatenation. We next tested how NHEJ affects split transgene reconstitution by pharmacologically inhibiting DNA-PK with the drug Nu7441 (Figure 5D-F). We observed a significant reduction in dual vector (Figure 5D-E) or single vector transduction (Figure 5F) under DNA-PK inhibition. This suggests NHEJ positively regulates ITR-ITR joining.

### Blocking HDR increases NHEJ-driven concatenation

Considering the opposing effects on split transgene reconstitution we observed when inhibiting these two main pathways of DSB repair, we hypothesized that after uncoating, HDR factors are preferentially recruited to VGs but are less efficient at concatenation than NHEJ. This suggests that blocking HDR allows NHEJ-mediated concatenation to predominate and thus there is an increase in ITR-ITR joining. To test this hypothesis, cells were pretreated with B02 or DMSO then transduced U2-OS^LacI-mNG^ cells with rAAV2/2.lacO.64 array vectors, fixed at 48 hpt and immunostained for Rad51 (HDR) and 53BP1 (NHEJ). High-content imaging revealed Rad51 colocalization with nearly 100% of VG foci imaged in DMSO treated cells, and a dramatic loss of this colocalization in B02 treated cells (Figure 5G). Concomitantly, B02 treatment increased 53BP1 colocalization from ∼10% to >60% of VGs (Figure 5G).

## DISCUSSION

Effectively overcoming the packaging size constraint would resolve a major limitation for gene delivery with rAAV. The inefficiency of trans-splicing dual vectors has been partially solved by split inteins to circumvent the protein coding limitation, but these still employ strong ubiquitous promoters that offer little control over expression^38^. Elements such as tissue-specific promoters, inducible promoters, enhancer arrays, UTRs/miRMA binding cassettes and other strategies for fine-tuning expression levels and spatiotemporal control must be in *cis* with the ORF they are controlling. Leveraging these approaches to control gene expression therefore necessitates concatenation of multiple VGs. In addition to payload expansion, concatenation is presumed to be an important mechanism for persistence of rAAV delivered DNA and transgene expression^39^.

Significantly advancing rAAVs as versatile gene delivery vectors will require a better understanding of the mechanistic underpinnings of rAAV biology. By investigating mechanisms of concatenation and expression from the point of view of the host cell, here we identify novel druggable targets for increasing dual vector transduction. Taken together, our results propose a model for host-vector interactions and pathway decisions that govern concatenation and expression of dual vectors (Figure 6). Shortly after uncoating, dsDNA VGs associate with histones and are flagged by the host cell by marks that are the same as those used for a double-strand break (Figure 6A). HDR factors such as Rad51 and BRCA1 are preferentially recruited to VGs, which blocks NHEJ factor recruitment, and HDR is the slower concatenation mechanism (Figure 6B). Blocking HDR allows the more efficient NHEJ to concatenate VGs, resulting in an increase in split transgene reconstitution (Figure 6C). Under transient Rad51 inhibition, we achieve dual vector transduction levels comparable to 100-fold higher vector doses without drug. Lowering vector doses is a key step toward ensuring safety of gene therapies. Our results also suggest that diseases of deregulated HDR, such as BRCA deficient cancers, may potentially confer an advantage for rAAV clinical applications in some settings.

**Figure 6.**
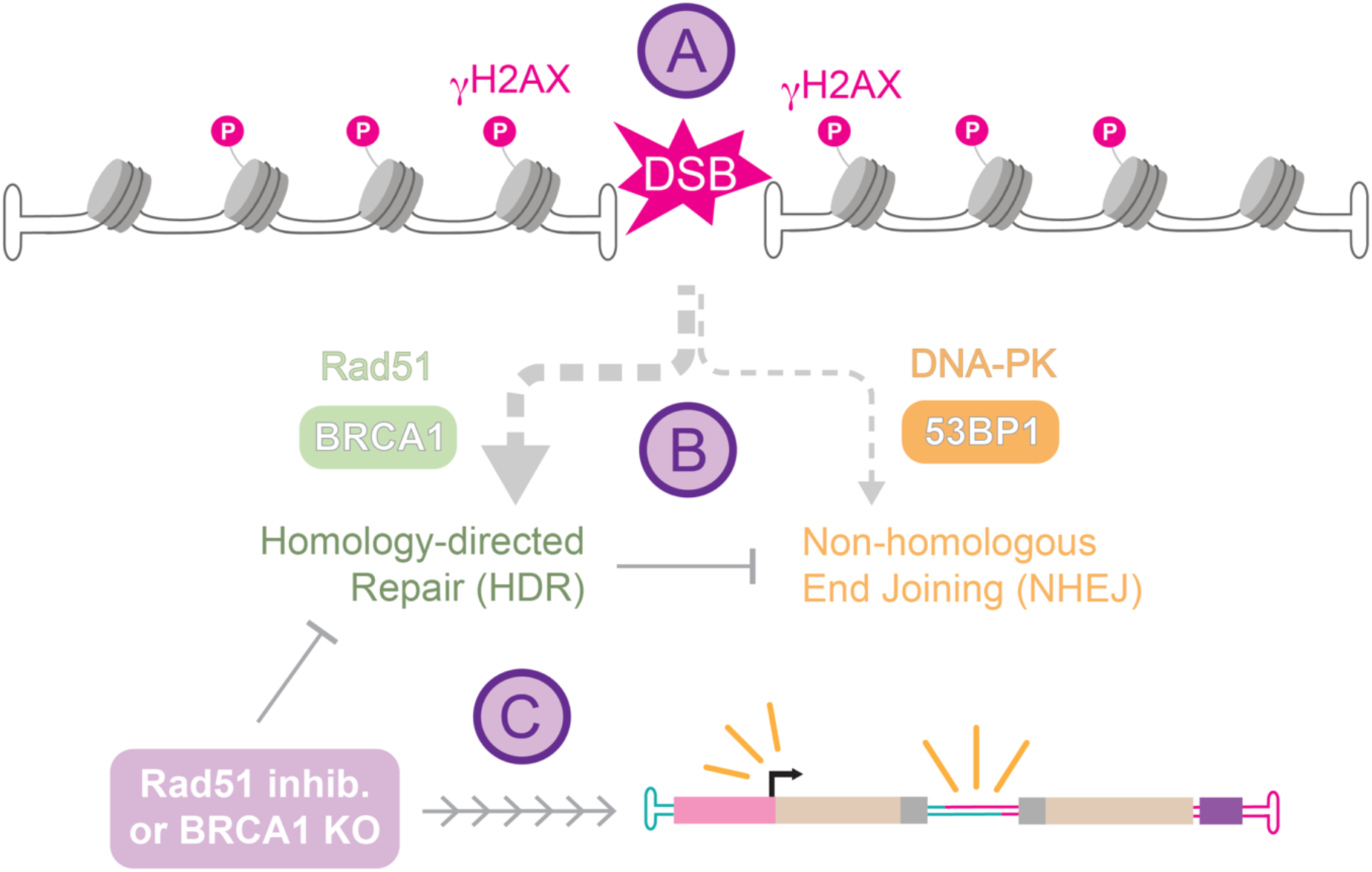
Model for processing of rAAV VGs by cellular DNA repair pathways. **(A)** Early in transduction, rAAV vector genomes quickly associate with histones and are marked as double-strand breaks (DSB) by the host cell as evidenced by staining for γH2AX. **(B)** HDR machinery appears to be recruited first (as detected with antibodies to BRCA1 and Rad51), which may block successful recruitment of NHEJ factors (such as DNA-PK and 53BP1). **(C)** Inhibition or loss of HDR machinery promotes increased expression and concatenation, which synergize to increase dual vector transduction.

It is widely believed that the AP fragment in HB dual vectors improves efficiency by promoting HR/HDR between VGs^26,27,40^, however this has not been formally tested in this context. Here, we present evidence that HR/HDR inhibits split transgene reconstitution, regardless of AP inclusion. This suggests that AP does not primarily function through HR/HDR. We observed similar effects on TS or HB dual vector transduction when we inhibit HDR or NHEJ (Figure 5), suggesting that these DSB repair pathways function similarly in ITR-ITR joining or AP overlapping. Upstream of both HDR and NHEJ, the MRN complex recognizes DSB free ends, and has been implicated as inhibitory to wt AAV infection/replication and to rAAV transduction, including rAAV VG foci formation^15–17,35^. The MRN complex may function in multiple ways in response to rAAV VGs, as it does in the response to cellular DSBs, and this may explain why we did not observe enrichment of any MRN components in our genome-wide screen using the HB dual vectors.

The novel vector tools introduced in this study enable us to investigate mechanisms of transduction at individual VG resolution and therefore open new avenues for future studies. Additional research is needed to clarify exactly how Rad51 and HDR impede concatenation of VGs. The canonical role for Rad51 in HR/HDR is to coat ssDNA after resection at the DSB end and promote homology search and strand invasion^41^. One possibility is that the majority of ITRs are in the “T” conformation and are technically dsDNA, and not coated by Rad51. Such ITRs cannot perform efficient homology searches for neighboring ITRs with which they can concatenate. Perhaps this dead-end Rad51 binding impedes NHEJ factor recruitment and by extension concatenation through non-HDR pathways. It is also unclear specifically how Rad51 inhibition increases the proportion of transcriptionally active VGs. Highly transcribed genes are known sites of genomic instability, and how the cell balances expression and repair is an actively researched topic^42,43^. One possibility is that Rad51 is involved in the cellular response to R-loops formed at the strong CMV promoter in our novel LacO.MS2 array reporter VGs. This rAAV vector system therefore presents a new tool for studying fundamental mechanisms of gene regulation and DNA repair.

The LacI-mNeonGreen expressing cell lines have excellent signal to background ratio at VG foci due to low level expression of LacI-mNG. While this enables detection of very small foci early in transduction and robust image analysis at all timepoints, it unfortunately limits our ability to quantify terminal sizes of episomes reliably by imaging. The pool of free LacI-mNG in the nucleus becomes depleted as transduction progresses, and it is unclear which, if any, VG arrays are saturated; only saturated VGs can be used to quantify size. Future studies may incorporate clonal lines with mid-range expression levels to enable size quantification under DDR or other pathway perturbations.

Chromatinization and epigenetic regulation are just beginning to be appreciated by the field as major determinants of gene therapy outcomes^44,45^. Epigenetic regulation is a well-known orchestrator of mammalian gene expression or silencing, as well as genome maintenance and repair, but has been only minimally examined in the context of AAV viral and vector genomes contexts. Active^46,47^ and repressive^47–49^ histone marks have been observed on VGs at late timepoints in transduced cultured cells and in mouse and NHP models. In NHPs, VGs extracted years post injection exist episomally as monomers and high molecular weight concatemers with nucleosomal structure^50^. It is unclear which VGs are expressing versus silenced, but moreover, how and when epigenetic control is exerted on VGs remains understudied. Here, we provide evidence for VG chromatinization very early in transduction in cell culture, and suggest a previously underappreciated role for epigenetic regulation in coordinating concatenation through DSB repair pathways. A limitation of our approach is that it cannot distinguish whether VGs are recruited to host genomic sites of active DNA damage repair and then hijack it, or whether VGs independently recruit these factors. The former has been suggested as the mechanism for wt AAV viral infections^32^, but the viral Rep proteins implicated in these studies are absent in the recombinant AAV setting.

rAAV VGs remain predominantly episomal, but integrations can occur at low levels and are of concern for clinical applications. While the present study does not examine integration of VGs, the effects of HDR/NHEJ manipulation on integration are an exciting avenue for future study, particularly in the context of rAAV-delivered HDR donor templates for CRISPR-mediated editing or knock-in strategies. The tools introduced in this study will be useful for studying vector-host interactions driving many aspects of rAAV biology as well as fundamental cellular and molecular biology.

## MATERIALS AND METHODS

### Plasmids

*Trans* rep-cap pAAV2/2 (#104963), *cis* pAAV.CMV.Luc.IRES.EGFP.SV40 (#105533) and Helper plasmids pAdDeltaF6 (#112867) were obtained from Addgene. ITR2.LacO.64 array *cis* plasmids were cloned from a pENTR 16.4 LacOR Tjian Lab vector (array of 16 lacO sites concatenated 4 times) into the ITR2-containing backbone from pAAV.CMV.EGFP.Luc. To create ITR2.LacO.64.CMV.mScarlet, The CMV promoter and mScarlet cassette were PCR amplified adding a peroxisome targeting sequence (SKL) to the C-terminus; the amplicon was ligated into RE digested ITR2.lacO.64. ITR2.LacO.64.CMV.MS2 was generated from RE digest to remove mScarlet from ITR2.LacO.64.CMV.mScarlet and RE digest of pMBSV5 (Singer Lab), to yield a fragment containing 24 repeats of the MS2 stem-loop sequence, which was ligated into the ITR2.LacO.64.CMV backbone. All plasmid stocks were propagated in StbL2 competent cells, with some intermediate cloning steps carried out in DH5alpha.

### Vector Production and Purification

Vectors were produced by triple-plasmid transfection in HEK293T cells as previously described^51^. For large-scale production and purification, at least ten 15cm dishes of HEK293T were transfected with the appropriate plasmids and cells + media were collected 72h later and subjected to 3 freeze/thaw cycles. Lysate was clarified by centrifugation, Benzonase treated for 30 min at 37C, then lysate was 22u filtered and purified by AVB Sepharose Hi-Trap prepacked columns following the manufacturer’s instructions on an AKTA pure FPLC system. All titers were assayed by taqman qPCR using primers and probes within the transgene cassette, or in the case of LacO.64 which consists only of the repeat array, the ITRs.

### Cell lines

HEK293T cells used for vector production were revived from a tube cryopreserved in 1984. To generate U2-OS^LacI-mNeonGreen^, U2-OS cells (UC Berkeley BDS Cell Culture Facility) cultured in DMEM (1 g/L glucose, 10% FBS) were transfected with a modified PiggyBac Transposase plasmid (System Biosciences) and a PGK.mNeonGreen-LacI.NLS.IRES.Puro integration plasmid using Fugene6 transfection reagent. After 48h, cells were subjected to Puromycin selection (1ug/mL) for 96h. Survivors were expanded for an additional 96h at maintenance (0.2ug/L) Puromycin concentration. Individual cells were then plated in 96 well plates using a FACS Aria. After 18 days, individual colonies were expanded to larger wells and to replicate CellCarrier Ultra Plates (Perkin Elmer) for initial screening on an OperaPhenix high-content imager. Live colonies were imaged with a 488 filter set, and those expressing mNeonGreen within an ideal threshold (nuclear signal between 200-2000) were passaged to replicate Falcon and CellCarrier Ultra plates for additional screening (48 clones total). Chosen colonies were then transduced with 1e9 vg/well rAAV2/2.lacO.64 crude vector preparation and live imaged at 48 hpt. The three colonies with best signal to background ratio at LacI-mNG foci were expanded and used for further experiments. To generate U2-OS^LacI-mNeionGreen.MCP-mCherry^ cells, one clonal line was transfected with PiggyBac Transposase and a UBC.tandemMCP2-mCherry plasmid with a neomycin resistance marker integration plasmid. Selection under 1mg/mL G418 was carried out as above, and clonal expansion and screening performed as above, imaged with mCherry filter sets after transduction with rAAV2/2. lacO.64.CMV.MS2.24.

### LacZ-Activated Fluorescence Assay (LAFA)

U2OS cells were seeded in DMEM (1 g/L glucose and 10% FBS) at a density of 2,500 cells/well in a 96-well plate. The following day, cells were pretreated with the pharmacological inhibitors B02 (5M or 10M) or Nu7441 (1.75M or 2.75M) in DMEM (high glucose and 10% FBS) for 4-16h. Spent media was replaced with DMEM containing the appropriate dose of unsplit or split LacZ vector, and B02 (5M or 10M), Nu7441 (1.75M or 2.75M), or DMSO, and 5% FBS. Cells were transduced for a total of 48 hours without further media changes. Following transduction, cells were washed with 1x PBS and lysed in plate using 50uL cold lysis buffer (2mM DTT, 25mM Tris-phosphate (pH 7.8), 2mM 1,2-diaminocyclohexane-N,N,Ń,Ń-tetraacetic acid, 1.25mg/ml lysozyme, 2.5mg/ml BSA, 10% glycerol, 1% Triton® X-100). To facilitate lysis, cells underwent two freeze/thaw cycles at -80 C and 37 C, respectively. 4-methylumbelliferyl-[beta]-D-galactopyranoside (MUG, 10mM in DMSO) was diluted 1:10 in LAFA buffer (100mM Sodium Phosphate Buffer (pH 7.0), 1mM MgCl2, 10mM 2-mercaptoethanol, 0.1% Triton® X-100) to a final concentration of 1mM. In a 96-well plate, 20uL of the cell lysate was added to 100uL of the MUG-LAFA buffer mixture, protected from light, and gently rotated at room temperature for 90 minutes. Fluorescence signal was read at 365nm/460nm (Ex/Em) using a Tecan Infinite® M1000. To account for any growth defects due to drug treatments, cell viability was quantified using a CellTiter-Glo® kit according to the manufacturer’s protocol, and this was used to normalize per well fluorescence.

### Screen and pathway-enrichment analysis

A lentiviral Brunello/Cas9 library was purchased from the Broad Institute Genetic Perturbation Platform (GPP) and used to infect 400 M U2-OS cells at MOI of 0.5. Puromycin selection began 48h later and continued for 96h. The surviving cell library was passaged once into 15cm plates and 40M cells were transduced with 5e3 vg/cell rAAV2/2.HB L and R vectors in biological duplicate. At 48 hpt, cells were collected and stained with MarkerGene™ according to the manufacturer’s protocol, sorted on a FACS Aria (BD), and gDNA extracted from each population with the Qiagen DNeasy Blood and Tissue kit according to manufacturers protocol. NGS and was performed at the Broad GPP as previously described^52^. Log-normalized sgRNA barcode counts from FACS-sorted MUG-negative cells were subtracted from FACS-sorted MUG-positive cells and used to model a hypergeometric distribution using the Broad GPP Web Portal. Genes with fewer than 2 unambiguous sgRNA perturbations were culled from this analysis. The most significant 5% of positive regulators (i.e., genes that, when perturbed, have increased dual vector transduction) were submitted as a gene list for Reactome pathway analysis^53^.

### High-content imaging and analysis

Cells were plated in CellCarrier Ultra 96 well plates (Perkin Elmer) and transduced/treated with the appropriate drugs and vectors. For live cell imaging (Movie 1) cells received 1e6 vg/cell of rAAV2/2.lacO.64. LacI-mNG was visualized with a 488 filter set on an Opera Phenix automated imager at 37C and 5% CO2 from 2-24 hpt and from 26-48 hpt, at 1 frame every 8 minutes. Images were combined in imageJ for Movie 1. For high-content imaging experiments, cells were fixed and stained with Hoechst to facilitate nuclear segmentation, and immunostained where indicated with the below primary antibodies overnight at 4C for detection by an Alexa647-conjugated secondary antibody. ROIs comprising up to 50% of the well surface were imaged with 11 z-planes at 0.5 micron steps. MIPs were processed and quantified using Harmony image analysis software using an appropriate analysis pipeline (Tables 1-3). Well and object results were imported into Graphpad Prism for data visualization.

### Antibodies and IF dilutions

53BP1 – Abcam ab175188, 1:10,000; Phospho Histone H2A.X – Cell Signalling Technology 25775, 1:2500; BRCA1 Santa Cruz sc-6954 1:2000; Rad51 – Abcam ab133534, 1:1000.

## Acknowledgements

We thank Eva Andres-Mateos, Heikki Turunen, Mohammadsharif Tabebordbar, Thomas Graham, Claudia Cattoglio, Erin Merkel, Nam Che, John Doench, Eric Zinn, Dirk Hockemeyer, Antonio Maffia, and Tyler Huycke for helpful discussions. Thanks to Mathieu Nonnenmacher for critical reading of the manuscript. We thank Rob Singer and Luk Vandenberghe for plasmids. Thanks to the Genetic Perturbation Platform (Broad Institute) for assistance with processing screening samples. We thank Mary West of the High-Throughput Screening Facility (HTSF) at UC Berkeley. This work was performed in part in the HTSF, that provided the OperaPhenix high content imager. Funding was provided by the National Institute of Health grants (1-U54CA231641-01), the California Institute for Regenerative Medicine Training Program EDUC4-12790, and the Howard Hughes Medical Institute (34430, R. T.).

